# BDH1-Dependent Ketone Body Metabolism Maintains Müller Cell Homeostasis and Retinal Function

**DOI:** 10.64898/2025.12.27.696636

**Authors:** Richa Garg, Eshani Karmakar, David DeBruin, Shauna Prasad, Ethan Naquin, Michelle Brennan, Niloofar Piri, Oleg Kisselev, Jaya P. Gnana-Prakasam

## Abstract

Ketone body metabolism serves as an auxiliary regulator of cellular energetics and redox balance, particularly during prolonged fasting and carbohydrate restriction, yet its role in retinal homeostasis under physiological conditions remains poorly defined. β-hydroxybutyrate dehydrogenase 1 (BDH1) is a mitochondrial enzyme that interconverts acetoacetate and β-hydroxybutyrate, and is required for efficient ketone utilization. Here, we investigated the impact of impaired endogenous ketone metabolism on retinal function using global and retinal pigment epithelium (RPE)–specific BDH1 knockout (KO) mice. Global BDH1 KO mice showed reduced circulating β-hydroxybutyrate and blunted fasting-induced ketone elevations, accompanied by ganglion cell loss, structural abnormalities on fundus and OCT imaging, and diminished scotopic and photopic electroretinogram (ERG) a- and b-wave amplitudes, consistent with impaired photoreceptor responses and downstream bipolar and Müller cell signaling. In contrast, RPE-specific BDH1 KO mice exhibited no changes in ERG responses or retinal morphology. Transcriptomic and molecular analyses in global KO retinas revealed disrupted Müller cell homeostasis, including reduced CAMKII-CREB activation, which is required for EAAT1 glutamate transporter expression. Administration of exogenous β-hydroxybutyrate, *in vitro* and *in vivo,* restored CAMKII–CREB–EAAT1 signaling, glutamate uptake, and antioxidant gene expression in BDH1 KO mice, demonstrating a central role for ketone bodies in Müller cell metabolic support, glutamate homeostasis, and redox balance. Together with reduced BDH1 expression in human AMD retinas, these findings identify the BDH1–β-hydroxybutyrate axis as a critical metabolic pathway for Müller cell function and retinal integrity, and highlight ketone metabolism as a potential therapeutic target in degenerative retinal diseases.

## Introduction

Ketone bodies (KBs) have emerged as critical metabolic intermediates that function not only as alternative energy substrates during limited glucose availability, but also as signaling molecules regulating cellular metabolism, inflammation, and redox homeostasis (1). Under physiological conditions, circulating KB concentrations range from ∼0.05–0.4 mM and increase to 1-2mM during prolonged fasting, carbohydrate restriction, or poorly controlled diabetes (2). The liver is the principal site of ketogenesis, producing the ketone bodies, β-hydroxybutyrate (BHB), acetoacetate (AcAc), and acetone. However, extrahepatic tissues including intestine, kidney, brain, and retina also produce ketones locally (3).

Ketogenesis is initiated by mitochondrial β-oxidation of fatty acids, generating acetyl-CoA, which is converted to AcAc by mitochondrial enzymes, 3-hydroxy-3-methylglutaryl-CoA synthase 2 (HMGCS2) and HMG-CoA lyase (HMGCL). 3-hydroxybutyrate dehydrogenase-1 (BDH1) then catalyzes the conversion of AcAc to BHB, with BHB comprising approximately 70% of circulating KBs (4, 5). Although the liver synthesizes ketones, it lacks enzymes required for ketone oxidation. Extrahepatic tissues instead utilize BHB as fuel to meet energy demands during periods of limited glucose availability. KB oxidation begins when BDH1 converts BHB back to AcAc, followed by reconversion to acetyl-CoA for entry into the tricarboxylic acid (TCA) cycle and ATP production (6). BDH1 is a unique enzyme that catalyzes the reversible interconversion between AcAc and BHB, linking ketone production and utilization.

The retina is one of the most metabolically demanding tissues in the body. Selective expression of the rate-limiting ketogenic enzyme HMGCS2 in the retinal pigment epithelium (RPE), together with ketolytic enzyme expression in photoreceptors and other retinal cell types, identifies the RPE and neural retina as key sites for ketogenesis and ketolysis respectively. In addition to uptake of hepatic-derived BHB through monocarboxylate transporters (MCTs), RPE cells in the retina produce BHB locally (7). Notably, RPE cells phagocytose the shed photoreceptor outer segments (POS) enriched in fatty acids, fueling mitochondrial ketogenesis and BHB production, which can be shuttled to adjacent retinal neurons as a metabolic substrate (7, 8).

BDH1 plays a central role in both BHB synthesis and utilization. Mouse models lacking BDH1 exhibit reduced circulating BHB and increased hepatic lipid accumulation during fasting (9). Conversely, BDH1 overexpression mitigates oxidative stress, inflammation, and apoptosis in mouse models of fatty liver disease and diabetic nephropathy (10, 11). BHB exerts anti-inflammatory effects through inhibition of the NLRP3 inflammasome, a critical mediator of proinflammatory cytokines such as IL-1β and IL-18 (12, 13). In streptozotocin (STZ)-induced diabetic mice, systemic BHB treatment reduced the expression of NLRP3 inflammasome components (NLRP3, ASC, caspase-1) and proinflammatory cytokines in the retina, accompanied by decreased apoptosis in the outer nuclear layer (ONL) (14). Furthermore, BHB treatment enhanced the retinal levels of brain-derived neurotrophic factor (BDNF) and connexin 43 (Cnx43), both of which are downregulated in diabetic retinopathy, contributing to cellular apoptosis (14, 15). BHB also activates the hydroxycarboxylic acid receptor 2 (HCAR2/GPR109A), which plays a crucial role in anti-inflammatory and neuroprotective processes, such as preserving the blood-retinal barrier integrity, an essential defense against vascular leakage in retinal degenerative diseases (16). In addition, exogenous BHB activates nuclear factor erythroid 2–related factor 2 (Nrf2), a transcription factor that induces antioxidant genes such as HO-1 and NQO1 (17), thereby mitigating oxidative stress-induced retinal injury. In vitro studies further demonstrate that BHB protects ARPE-19 cells from high glucose-induced cellular damage (18). Moreover, in the rd10 mouse model of retinal degeneration, ketogenic diet delayed retinal degeneration and improved electroretinogram (ERG) responses (19). Collectively, these findings highlight the therapeutic potential of exogenous BHB and ketogenic diets in alleviating oxidative stress, inflammation, and neurodegeneration in animal models of retinal degeneration. Despite these advances, the role of constitutive endogenous ketone metabolism in maintaining retinal homeostasis remains unclear.

Accumulating evidence indicates that ketogenic interventions exert neuroprotective effects across diverse neurodegenerative disorders (20). Given shared pathogenic mechanisms between neurodegenerative brain diseases and retinal disorders such as age-related macular degeneration (AMD) and chronic optic neuropathies, impaired endogenous ketone metabolism may be an underrecognized critical determinant of retinal dysfunction. Here, we investigated the role of constitutive BHB in maintaining retinal structure and function using global and RPE-specific BDH1 knockout mouse models. Our findings reveal that BDH1 loss compromises Müller glial metabolic support, disrupts glutamate handling, increases oxidative stress and accelerates neurodegeneration.

## Methods

### Animals

BDH1 knockout (KO) mice were generated commercially through Cyagen Biosciences (Santa Clara, CA, USA) using CRISPR/Cas9-mediated genome editing. The mouse *Bdh1* gene, located on chromosome 16 and comprised of seven exons, was disrupted by deleting exons 3-7 using a pair of guide RNAs (gRNAs). Cas9 mRNA and gRNAs targeting the *Bdh1* locus were synthesized by in vitro transcription and co-injected into fertilized mouse zygotes to generate KO founder animals. Offspring were genotyped by PCR and confirmed by Sanger sequencing. Verified founder KOs were crossed with wild-type (WT) mice to assess germline transmission and produce F1 heterozygotes, which were subsequently intercrossed to obtain homozygous KO, heterozygous, and WT littermates on a C57BL/6J background for experiments. All genotypes were independently validated by Transnetyx Inc. (Cordova, TN, USA). RPE-specific BDH1 conditional KO mice were generated by crossing *Bdh1* floxed mice (*Bdh1 ^fl/fl^*), kindly provided by Dr. Daniel P. Kelly at the University of Pennsylvania (21), with BEST1-Cre mice (Stock # 017557, The Jackson Laboratory, Bar Harbor, ME, USA), generously provided by Dr. Joshua Dunaief at the University of Pennsylvania (22) to obtain *Cre^+^Bdh1^fl/fl^* mice. All mouse strains were confirmed negative for rd mutations prior to experimental use. Mice were housed in animal facility at the Saint Louis University School of Medicine under controlled temperature and humidity with a 12h light/dark cycle and ad libitum access to food and water. Age and sex-matched mice were used for all experiments. All animal procedures were approved by the Saint Louis University Institutional Animal Care and Use Committee and adhered to the Association for Research in Vision and Ophthalmology Statement for the Use of Animals in Ophthalmic and Vision Research.

### Postmortem Human Retina

We communicate with over 20 eye banks nationally to obtain human donor eyes with a diagnosis of AMD, as well as age-matched controls, for research purposes. Since these human donors were deidentified and ocular tissues were provided solely for research purposes, IRB approval was obtained by the tissue donor center and was not required by the local research institution. Donors with a history of other intraocular pathologies, such as glaucoma and retinal vascular diseases, including diabetic retinopathy, were excluded. Death to preservation time was less than 6 hours to minimize intracellular degeneration. After enucleation, whole globes were placed in either Davidson’s fixative or 10% formalin. Within 24 hours after fixation, whole globes were sectioned and the macula identified. Tissue sections were paraffin-embedded at the Digestive Disease Research Center (DDRC) in Washington University. Four-micrometer sections from the macular area were stained with hematoxylin and eosin (H&E) and histologically reviewed by two retina specialists to assess tissue quality and confirm the presence of drusen in AMD donor sections. Age-matched control slides were also evaluated to confirm that tissue quality was preserved, and that no macular pathology was present. When necessary, additional sectioning was performed to obtain sections as close to the center of the macula as possible. After histologic evaluation was completed, an additional consecutive unstained section was obtained for the current study.

### Immunostaining

Paraffin-embedded tissue sections (4–5 µm thick) were deparaffinized in xylene and subsequently rehydrated through a graded ethanol series (100%, 95%, 80%, 70%, and 50%), followed by rinsing in distilled water to remove residual solvents. Antigen retrieval was performed in 10 mM sodium citrate buffer (pH 6.0) by heating at 95–100 °C for 20 min, after which the slides were allowed to cool gradually to room temperature and rinsed thoroughly with phosphate-buffered saline (PBS). Non-specific antibody binding was minimized by incubating the sections with 3–5% bovine serum albumin (BSA) in PBS for 30–60 min at room temperature. The sections were then incubated overnight at 4 °C with primary antibodies diluted (1:50–1:200) in the same blocking buffer. Following three to four washes with PBST (PBS containing 0.05% Tween-20), the sections were incubated with Alexa Fluor conjugated secondary antibodies (1:400) for 1–2 h at room temperature in the dark. Nuclei were counterstained with DAPI for 5–7 min, followed by washing in PBS. The slides were then mounted using an anti-fade mounting medium. Fluorescent signals were visualized, and representative images were acquired using a Leica SP8 confocal microscope (Leica Microsystems, Wetzlar, Germany) under identical acquisition settings.

### Metabolic Measurements

Total serum ketone bodies were measured using the Autokit Total Ketone Body Assay (Wako Life Sciences Inc., Richmond, VA, Catalog # 415-73301 and # 411-73401). Serum β-hydroxybutyrate levels were determined using the LiquiColor reagent kit (StanBio, Boerne, TX, Catalog # 2440-058), and triglycerides were measured using the Infinity Triglyceride Kit (Thermo Fisher Scientific, Catalog # TR22421) according to the manufacturers’ protocols. Blood glucose levels were determined from tail vein samples using a Contour Next EZ glucometer (Ascensia Diabetes Care). For the glucose tolerance test (GTT), mice were fasted for 15h and injected intraperitoneally with D-glucose (2g/kg body weight). Blood glucose levels were measured at 0, 15, 30, 60, 90, and 120 min post-injection.

### Electroretinography (ERG)

Mice were dark-adapted overnight and anesthetized by subcutaneous injection of ketamine (80 mg/kg) and xylazine (15 mg/kg). Pupils were dilated with 1% atropine sulfate. During testing, a heating pad controlled by a rectal temperature probe maintained body temperature at 37–38 °C. Full-field flash ERGs were recorded using a UTAS BigShot apparatus (LKC Technologies) and corneal cup electrodes, as previously described (23). The reference electrode needle was inserted subcutaneously at the skull. Rod and cone visual function were measured using full-field scotopic and photopic flash ERGs in a repeated measures design. Mice, in groups of five, were dark-adapted, after which scotopic ERGs were recorded. Test flashes of white light, ranging from 2.5x10^-5^ cd·s m^-2^ to 80 cd·s m^-2^, were applied under dark-adapted (scotopic) conditions. Responses from several trials were averaged, and the intervals between trials were adjusted so that responses did not decrease in amplitude over the series of trials for each step. The recorded responses were low-pass filtered at 500 Hz. Amplitudes of the a- and b-waves were measured using EMWin version 9.8.0 (LKC Technologies).

### *In vivo* Retinal Imaging

*In vivo* retinal imaging was performed using a Micron IV system (Phoenix Research Laboratories Inc., Pleasanton, CA, USA). Mice were anesthetized with a ketamine/xylazine mixture, and pupils were dilated with 1% atropine sulfate. A drop of methylcellulose was applied to each cornea to maintain corneal hydration. Fundus and optical coherence tomography (OCT) images were acquired using the corresponding modules of the Micron IV system (24). Quantitative analysis of retinal layer thickness was performed using Insight software (Phoenix Research Laboratories).

### Wholemount Retina Preparation and Staining

Whole-mount retinas were prepared from BDH1 WT and BDH1 KO mice as previously described (25). Briefly, whole eyes were fixed in 4% PFA for 2 hours at room temperature. Following fixation, the cornea and sclera were separated from the choroid-RPE-retina-lens complex. After lens removal, the retina was carefully dissected from the RPE-choroid complex, and all subsequent steps were performed on ice. Retinas were incubated in blocking solution (5% BSA in PBS) for 40 minutes, followed by staining with FITC-conjugated peanut agglutinin (PNA; 0.2mg/ml, Sigma) at 1:100 dilution overnight at 4°C to label cone photoreceptors. The following day, retinas were washed three times in PBS for 20 minutes each, then mounted on slides using antifade mounting medium. Fluorescence images were acquired using a Keyence BZ-X800 fluorescence microscope, and image processing and quantification were performed using ImageJ software.

### Hematoxylin & Eosin Staining and Retinal Ganglion Cell Quantification

Eyes from BDH1 WT and BDH1 KO mice were enucleated, fixed in 4% paraformaldehyde (PFA), and paraffin embedded. 4μm-thick retinal sections were stained with H&E. Images were acquired using a Keyence BZ-X800 microscope, and retinal ganglion cells (RGCs) per 100 μm were quantified using ImageJ software (26).

### RNA Sequencing

Total RNA was extracted from WT and KO tissues using the PureLink RNA Mini Kit (Thermo Fisher Scientific). RNA-seq library preparation was performed by GTAC@MGI (Washington University, St. Louis, MO, USA) using 500 ng to 1 µg of total RNA per sample. Ribosomal RNA was depleted using RiboErase (Kapa Biosystems) following an RNAse H-based protocol. mRNA was fragmented at 94 °C for 8 minutes in reverse transcriptase buffer, followed by cDNA synthesis using SuperScript III Reverse Transcriptase (Life Technologies) and random hexamer primers. The resulting cDNA was subjected to end-repair to generate blunt ends. A single adenine was subsequently added to the 3′ ends, facilitating ligation to Illumina sequencing adapters. Subsequently, ligated fragments were PCR-amplified and sequenced on an Illumina NovaSeq X Plus platform with 150-base paired-end reads.

Base-calling and demultiplexing were performed using Illumina’s bcl2fastq software, allowing one mismatch in the index read. Reads were then aligned to the Ensembl release 101 mouse primary assembly using STAR version 2.7.9a (27). Gene-level counts were obtained from uniquely aligned, unambiguous reads using feature Counts (Subread version 2.0.3) (28) and normalized using edgeR’s Trimmed Mean of M-values (TMM) method (29). Differential expression analysis was performed using the limma R/Bioconductor package with the voomWithQualityWeights function to model the mean-variance relationship. Genes with raw p-values ≤ 0.05 were considered significant (30). Functional enrichment was assessed using the gage package in R/Bioconductor, evaluating gene set enrichment across Gene Ontology (GO) terms, MSigDB gene sets, and KEGG pathways (31). The analysis tested whether log2 fold-changes within a given term significantly differed from the log2 fold-changes of genes outside the term. Cell-type deconvolution was performed using the Dtangle algorithm implemented in the Granulator R package (version 1.14.0). The RNA signature matrix was generated from the Mouse Retina Cell Atlas (MRCA) single-cell sequencing dataset, which profiles 138 distinct cell types (32). The Dtangle algorithm was applied to bulk RNA-seq data to estimate the relative abundance of major retinal cell types based on MRCA-derived gene signatures. For cell-type marker selection, marker genes were designated by integrating literature-based searches and established cell-type marker databases (32–34). Because some markers are shared across closely related cell types, particularly within neuronal or glial lineages, partial overlap among marker sets was expected and accounted for during interpretation (33).

### Quantitative Real-Time PCR (qPCR)

Total RNA from animal tissues and primary Müller cells were extracted using the PureLink RNA Mini Kit (Thermo Fisher Scientific) and quantified spectrophotometrically at 260nm. cDNA was synthesized from 100ng RNA using the High-Capacity cDNA Reverse Transcription Kit (ThermoFisher Scientific) according to the manufacturer’s protocol. The cDNA was diluted 1:10 and used for PCR to assess the gene expression on an ABI Quant Studio3 real-time PCR system with PowerUp^TM^ SYBR^TM^ Green Master Mix (Applied Biosystems, Waltham, MA, USA, Cat# A25742). Relative mRNA expression was determined using the ΔΔCt values normalized to 18S rRNA. Primer sequences used are listed in Table 1.

**Table 1:**
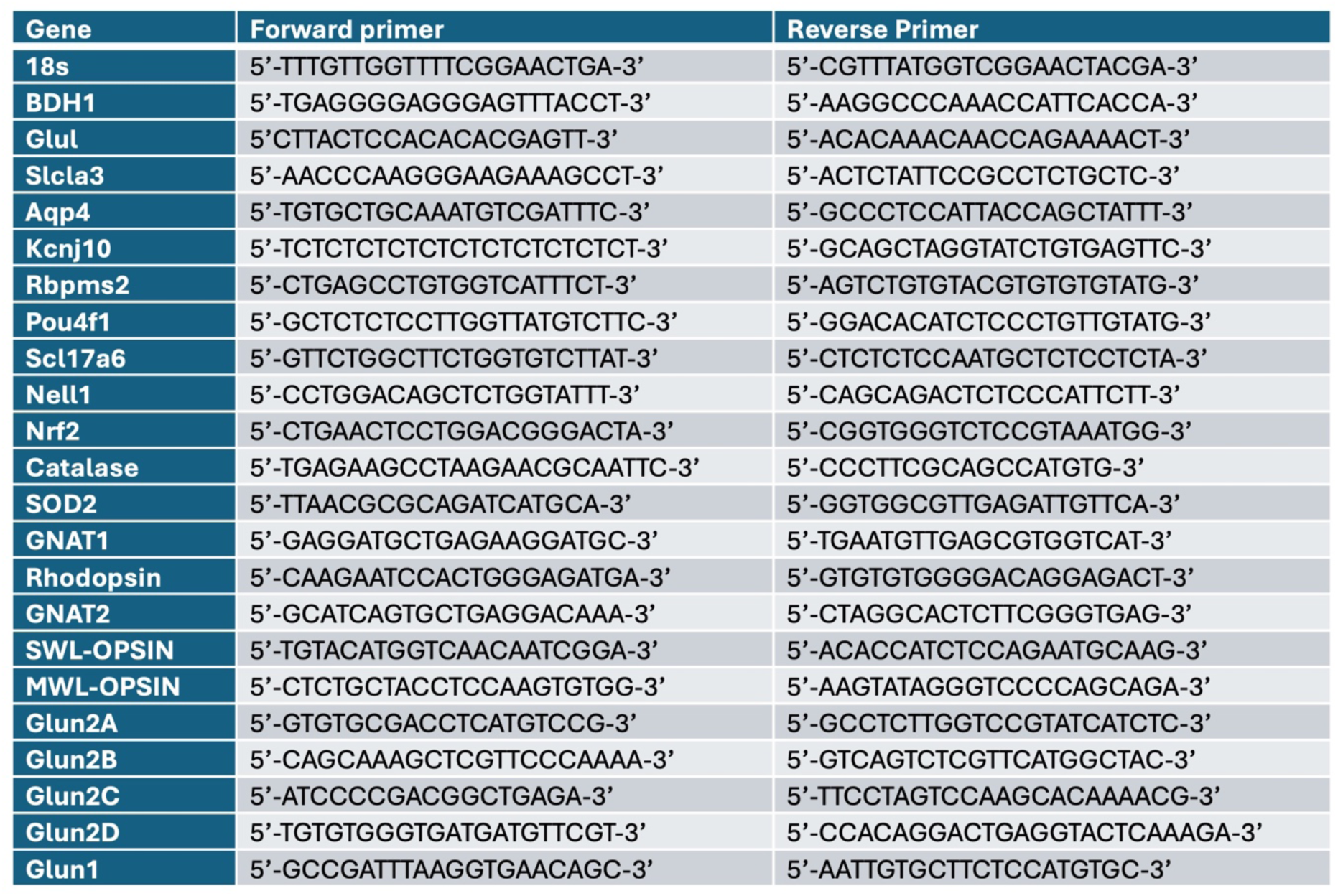
Primes used.

### Primary Mouse Müller Cell Culture

Primary Müller cells were isolated from 6 to 10-day-old pups as described previously (30). Cells were grown in DMEM (Gibco, Grand Island, NY, USA) supplemented with 10% fetal bovine serum (FBS) (Hyclone, Logan, UT, USA) and 5% penicillin/streptomycin (Invitrogen, Carlsbad, CA, USA) at 37 °C cell culture incubator with 5% CO_2_. Culture purity was confirmed as described previously by immunostaining for vimentin, a Müller cell marker (30, 35).

### Immunofluorescence

For primary Müller cells, 0.01×10^6^ cells/well were seeded on chambered slides and grown overnight in a cell culture incubator. Cells were fixed with cold 4% paraformaldehyde for 10 min and washed with PBS three times. Permeabilization was performed with 0.1% Triton X-100 for 5 min, followed by blocking with 3% bovine serum albumin (BSA) for 60 min. Cells were incubated with a primary antibody overnight at 4°C, washed, and then incubated with a fluorescence-conjugated secondary antibody for 90 min at room temperature and washed three times with PBS. Nuclei were counterstained with 0.25 μg/mL DAPI for 5 min. Coverslips were mounted using Prolong^TM^ Diamond Antifade Mountant (Thermo Fisher Scientific) and imaged using a Keyence BZ-X800 microscope.

### Calcium Assay

Intracellular calcium levels in primary Müller cells were quantified using the calcium-sensitive dye Fluo-4 AM (Catalog # F14201, ThermoFisher Scientific). The primary Müller cells, seeded in 96-well plates upon reaching confluence, were treated with BHB for 24 hours. Following treatment, cells were incubated with 5µM Fluo-4 AM and 5mM probenecid cocktail solution prepared in calcium and magnesium-free Hanks’ balanced salt solution (HBSS) for 1 h at 37°C. Fluorescence intensity was measured at excitation/emission wavelengths of 494/506nm using a BioTek microplate reader.

### Glutamate Uptake Assay

The glutamate uptake assay was performed as described previously (36). Primary Müller cells were treated with 5mM BHB for 24 hours and then incubated in calcium and magnesium-free HBSS for 30 minutes. Cells were subsequently exposed to HBSS containing calcium and magnesium supplemented with 100 μM glutamate for 3hours. The remaining glutamate concentration in the culture medium after the 3-hour incubation was determined using a colorimetric glutamate assay kit (SigmaAldrich; Catalog # MAK004-1KT) following the manufacturer’s protocol.

### Immunoblotting

Protein lysates from cells or tissues were extracted using RIPA Lysis buffer (ThermoFisher Scientific, Waltham, MA, USA, Cat# 89900) supplemented with DTT, PMSF, and protease/phosphatase inhibitor cocktail (Cell Signaling, Danvers, MA, USA, Cat# 5872S). Protein concentrations were quantified using the bicinchoninic acid (BCA) assay (Thermo Fisher Scientific, Waltham, MA, USA). Equal amounts of 20µg protein were separated by SDS-polyacrylamide gel electrophoresis, transferred to nitrocellulose membrane, and then blocked with 5% BSA for 1h. Membranes were incubated with primary antibodies overnight at 4°C, washed, and then incubated with HRP-conjugated secondary antibodies for 2h at room temperature. Blots were washed again, and the protein bands were visualized using an enhanced chemiluminescence (ECL) detection system (Thermo Fisher Scientific, Waltham, MA, USA). Antibody sources and dilutions used are listed in Table 2.

**Table 2:**
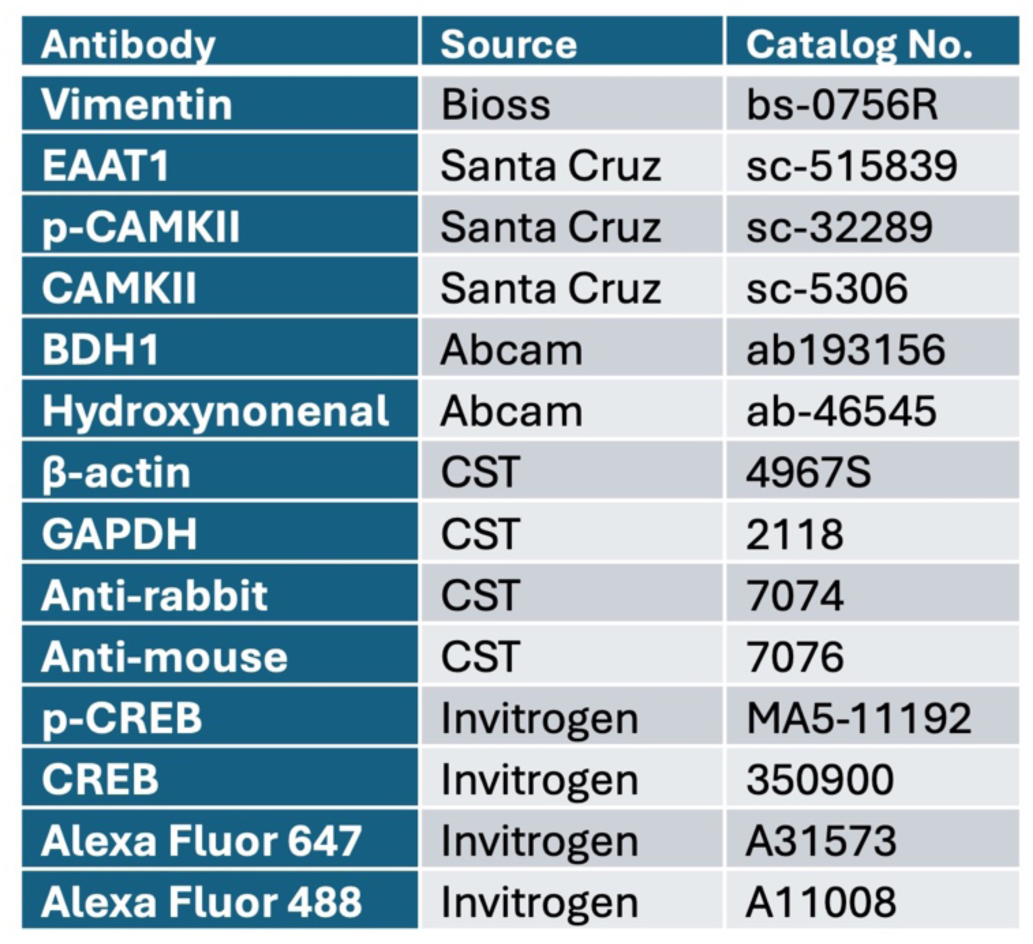
Antibodies used.

| Antibody | Source | Catalog No. |
| --- | --- | --- |
| Vimentin | Bioss | bs-0756R |
| EAAT1 | Santa Cruz | sc-515839 |
| p-CAMKII | Santa Cruz | sc-32289 |
| CAMKII | Santa Cruz | sc-5306 |
| BDH1 | Abcam | ab193156 |
| Hydroxynonenal | Abcam | ab-46545 |
| β-actin | CST | 4967S |
| GAPDH | CST | 2118 |
| Anti-rabbit | CST | 7074 |
| Anti-mouse | CST | 7076 |
| p-CREB | Invitrogen | MA5-11192 |
| CREB | Invitrogen | 350900 |
| Alexa Fluor 647 | Invitrogen | A31573 |
| Alexa Fluor 488 | Invitrogen | A11008 |

### Statistical analysis

Data are presented as mean ± SEM from three to four independent experiments, each performed in triplicate. For in vivo studies, age- and sex-matched mice (n=5-8 mice per group) were used. Fold changes were calculated by normalizing sample values to their respective controls. Statistical analyses were performed using GraphPad Prism10 software. Comparisons between two groups were analyzed using an unpaired two-tailed Student’s t-test, whereas multiple group comparisons were evaluated using one-way or two-way ANOVA followed by Tukey’s or Bonferroni post hoc tests, as appropriate. A p value < 0.05 was considered statistically significant.

### Data Availability

The datasets generated and analyzed during the current study are available from the corresponding author on reasonable request.

## Results

### BDH1 expression is diminished in human AMD retinas

AMD is a genetically predisposed, multifactorial, and progressive neurodegenerative disease of the retina characterized by oxidative stress, mitochondrial dysfunction, chronic inflammation, and disrupted metabolic homeostasis, ultimately leading to central vision loss in the elderly (37). Mitochondrial metabolic flexibility is essential for retinal integrity, and accumulating evidence implicates impaired energy metabolism as a critical driver of AMD pathogenesis (38, 39).

KBs, particularly ΒHB, exert neuroprotective effects by improving mitochondrial efficiency, attenuating oxidative stress, and modulating inflammatory pathways (12, 40). Circulating BHB levels decline with aging in mice (41). To determine whether alterations in ketone metabolism are associated with AMD, we examined BDH1 expression in postmortem human AMD retinal tissues. Immunohistochemical (IHC) analysis revealed abundant BDH1 immunoreactivity in the RPE, photoreceptors, inner nuclear layer (INL), inner plexiform layer (IPL) and ganglion cell layer (GCL) in healthy control donor eyes. In contrast, BDH1 staining was markedly reduced across multiple retinal layers in AMD samples (Figure 1A-B). Quantitative analysis confirmed a significant decrease in overall BDH1 signal intensity in AMD compared with age-matched controls, suggesting that impaired ketone metabolism may accompany AMD-related retinal degeneration. Given that ketone metabolism efficiency declines with age (41–43), loss of BDH1 expression may exacerbate metabolic inflexibility and oxidative stress in the aging retina. These findings underscore a potential role for BDH1 in AMD progression (Figure 1C), warranting further functional studies using BDH1-deficient models.

**Figure 1:**
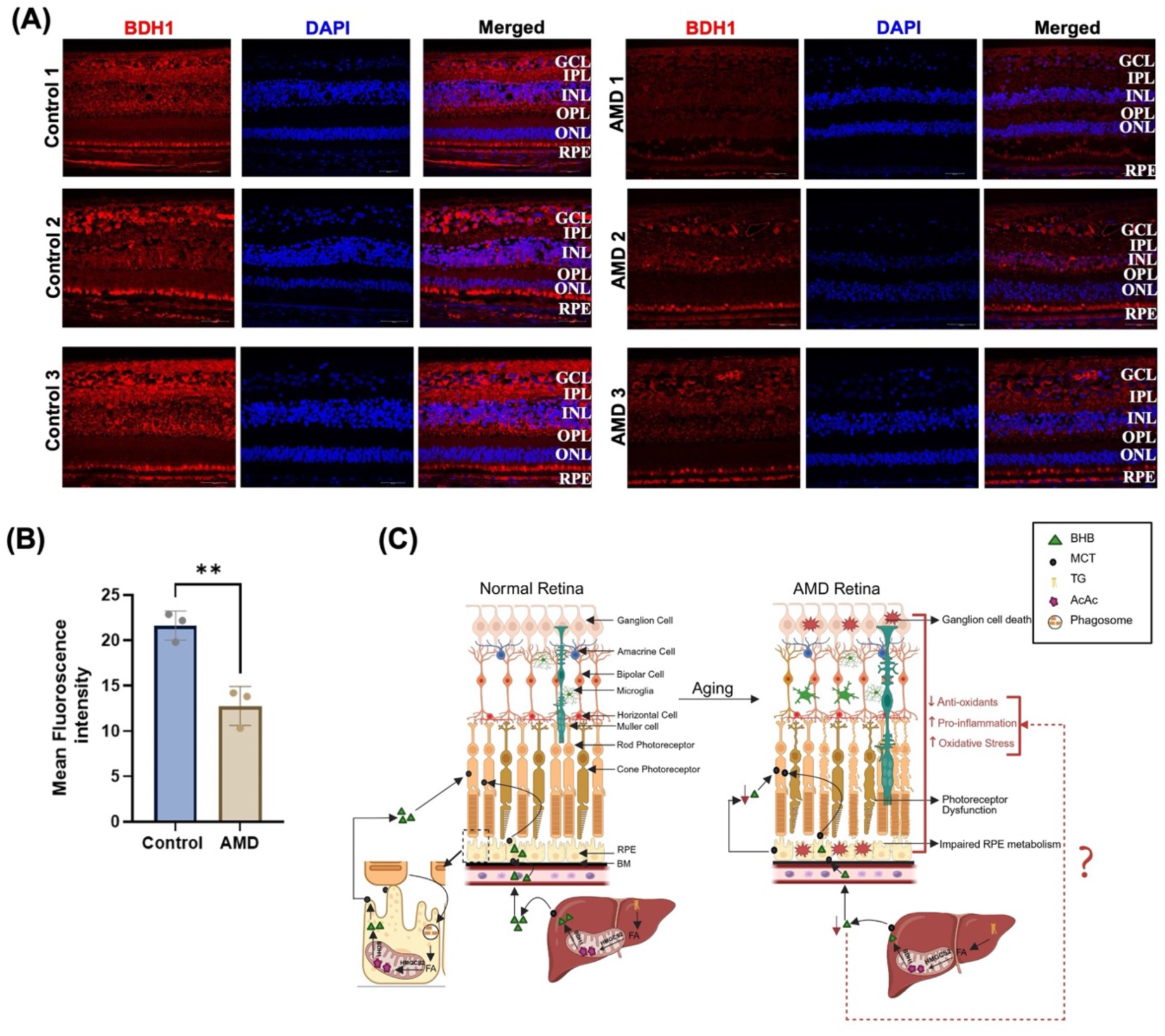
Reduced BDH1 expression in retinal tissues from patients with age-related macular degeneration (AMD). (A) Representative immunohistochemical images of postmortem human retinal sections demonstrating reduced BDH1 immunoreactivity across multiple retinal layers, including the RPE, photoreceptor cells, INL, IPL, and GCL in AMD donor samples compared with age-matched controls. (B) Quantification of BDH1 staining intensity confirms significant downregulation in AMD retinas. Scale bars, 50 µm. Data are presented as mean ± SEM; n = 3 per group; *P < 0.05 by unpaired *t*-test. (C) Schematic model illustrating the proposed impact of reduced BDH1-dependent ketone metabolism on retinal cellular homeostasis driving AMD pathology. Left panel is a transverse section of a healthy posterior eye highlighting major retinal layers, including the RPE, BM, photoreceptors (rods and cones), horizontal cells, bipolar cells, amacrine cells, ganglion cells, and resident microglia. In the healthy retina, the RPE phagocytoses photoreceptor outer segments and generates BHB, which is utilized by photoreceptors. Additionally, photoreceptors also take up liver-derived BHB via monocarboxylate transporters (MCTs). Right panel is a proposed aging-associated model in which reduced BDH1 expression and diminished BHB availability may impair metabolic support, increase oxidative stress, and enhance inflammatory signaling, promoting AMD-like pathology. Schematic created with BioRender. Abbreviations: Ganglion cell layer (GCL), inner plexiform layer (IPL), inner nuclear layer (INL), outer plexiform layer (OPL), outer nuclear layer (ONL), retinal pigment epithelium (RPE), Bruch’s membrane (BM), and triglycerides (TGs).

### Generation and systemic characterization of global BDH1 knockout mice

To examine the physiological role of BDH1 *in vivo,* we generated global BDH1 knockout (KO) mice using CRISPR/Cas9-mediated gene editing (Figure 2A). Quantitative RT-PCR and immunoblotting confirmed complete loss of BDH1 mRNA and protein in liver and ocular tissues of BDH1 KO mice (Figures 2B-C). Immunohistochemistry demonstrated strong BDH1 expression in WT mice across multiple retinal layers, including the RPE, photoreceptors inner segments, outer plexiform layer (OPL), inner plexiform layer (IPL), and GCL with no detectable staining in BDH1 KO retinas (Figure 2D), further confirming the absence of BDH1 in KO retinas.

**Figure 2:**
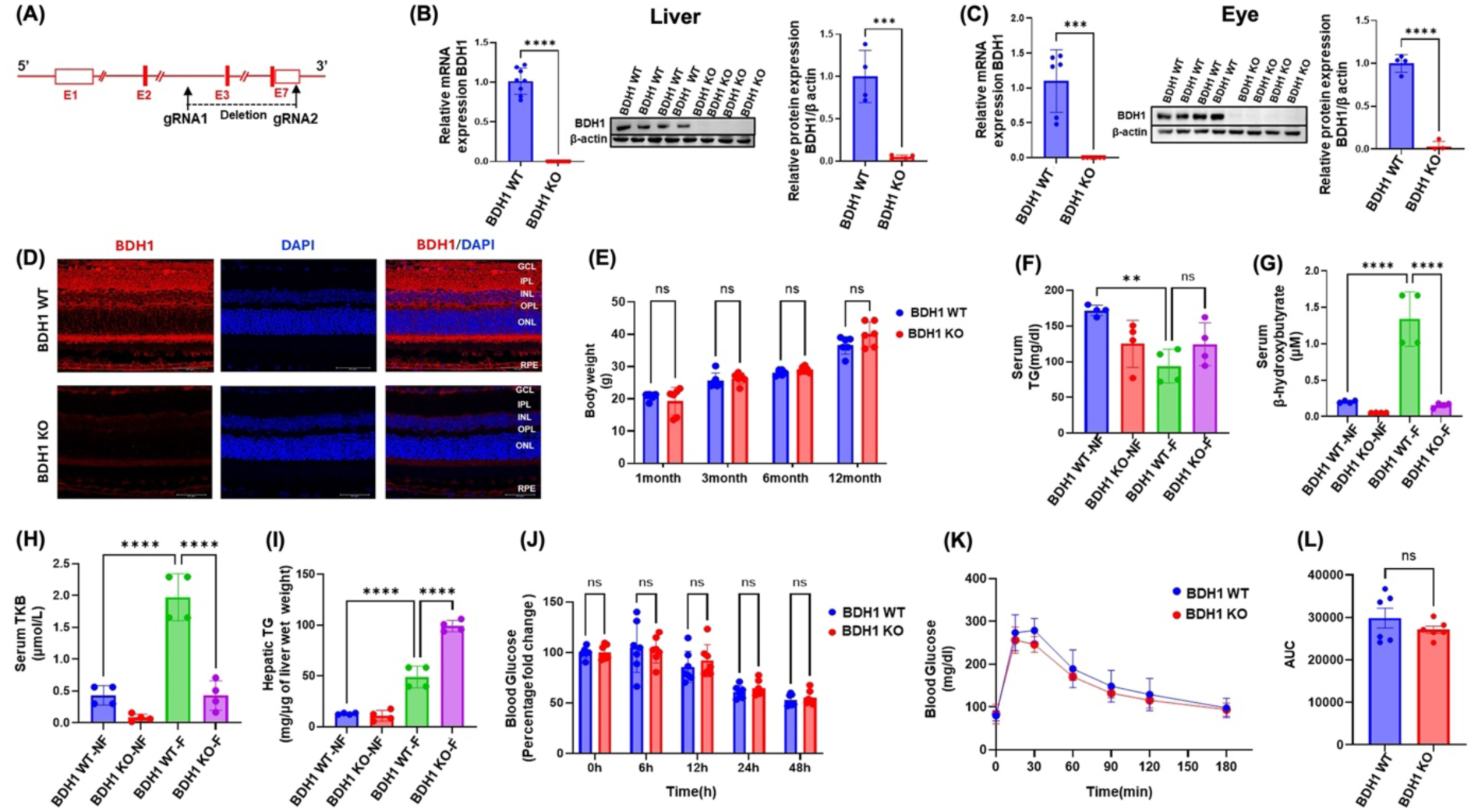
Generation and metabolic characterization of global BDH1 knockout mice. (A) Schematic of CRISPR/Cas9 strategy disrupting the *Bdh1* gene locus (total size, 30.15kb), deleting ∼14000bp region spanning exons 3-7 to generate the knockout mice. qRT-PCR and immunoblot analyses confirm complete loss of BDH1 mRNA and protein expression in (B) liver and (C) ocular tissues of BDH1 knockout (KO) mice. (D) Immunohistochemistry reveals robust BDH1 expression in WT retinas in the RPE, photoreceptor inner segments, IPL, OPL and GCL layers and absence of staining in BDH1 KO retinas. (E) Body weight at 1, 3, 6, and 12 months demonstrates normal growth in both groups. (F) Serum TG levels under fed and fasted conditions are comparable between genotypes. (G–H) Fasting markedly elevates circulating BHB and TKB levels in WT but not in BDH1 KO mice, indicating impaired ketogenesis. (I) Hepatic TG accumulation is increased in fasted BDH1 KO mice. (J) Blood glucose remains unchanged under fed and fasted conditions in both groups. (K–L) IPGTT and corresponding AUC analyses reveal normal glucose clearance in BDH1 KO mice. Data are mean ± SEM (n = 5–6 mice per group); *P≤0.05, **P≤0.001, ***P≤ 0.0001; ns, not significant. Abbreviations: outer plexiform layer (OPL), total ketone body (TKB), Intraperitoneal glucose-tolerance test (IPGTT), area-under-the-curve (AUC), non-fasted (NF), and Fasted (F).

To assess whether loss of BDH1 alters systemic metabolism, we monitored body weight and serum biochemical parameters. Body weights were comparable with no significant differences between the age- and sex-matched WT and BDH1 KO mice until 12months of age (Figure 2E), consistent with a previous report of normal growth and viability in BDH1-deficient mice (9). Under ad libitum feeding, serum triglyceride (TG) and glucose levels were comparable between the WT and KO mice (Figures 2F and J), indicating that basal systemic triglyceride and glucose levels remain intact in the absence of BDH1. As anticipated, fasting induced a significant increase in circulating BHB and total ketone bodies (TKB) in WT mice. In contrast, BDH1 KO mice on fasting showed a markedly attenuated rise in BHB and TKB (Figures 2G-H), accompanied by increased hepatic triglyceride accumulation (Figure 2I). These findings are consistent with previous reports demonstrating that impaired ketogenesis compromises hepatic lipid mobilization (1, 44). Notably, despite increased hepatic TG storage, circulating TG levels remained unchanged in WT and BDH1 KO mice under fasted conditions (Figures 2F and I).

There were no significant differences in fasting blood glucose levels between the WT and BDH1 KO mice at all measured time points from 0-48-hour fasting, indicating no glycemic impairment in BDH1 KO mice (Figure. 2J). Both groups exhibited comparable glucose clearance and area under the curve (AUC) during glucose tolerance test following intraperitoneal glucose challenge (Figure. 2K-L), suggesting that BDH1 deficiency selectively disrupts ketone metabolism without impairing systemic glucose levels or glucose tolerance. No sex-specific differences were observed across the measured parameters. Collectively, these findings demonstrate that while global BDH1 deficiency does not affect basal lipid or glucose metabolism, it profoundly impairs ketone production and fasting-induced hepatic triglyceride utilization, confirming the essential role of BDH1 in systemic ketone homeostasis.

### BDH1 deficiency leads to progressive retinal dysfunction

We next assessed the consequences of BDH1 loss on retinal function. Full-field ERG revealed significantly reduced scotopic (rod-mediated) and photopic (cone-mediated) responses in global BDH1 KO mice compared with age- and sex-matched WT controls (Figure 3A-B). 1-2 months old young BDH1 KO mice showed ∼20-25% reductions in scotopic and photopic b-wave amplitudes (Figure 3A and Suppl. Figure 1A respectively), indicating early dysfunction in bipolar cell signaling. There was an age-dependent decline, with 6-8 months old adult BDH1 KO mice exhibiting ∼37% reduction in a-wave and ∼35% reduction in b-wave amplitudes compared to age- and sex-matched WT mice, suggesting a progressive impairment in both photoreceptor and inner retinal function (Figure 3B). Consistently, there was a significant reduction in cone density and cone-specific gene expression in BDH1 KO compared to WT retinas (Suppl. Figure 1B-C; Suppl. Figure 1G-I). These findings indicate that BDH1 deficiency compromises visual signal transmission across multiple retinal cell layers.

**Figure 3:**
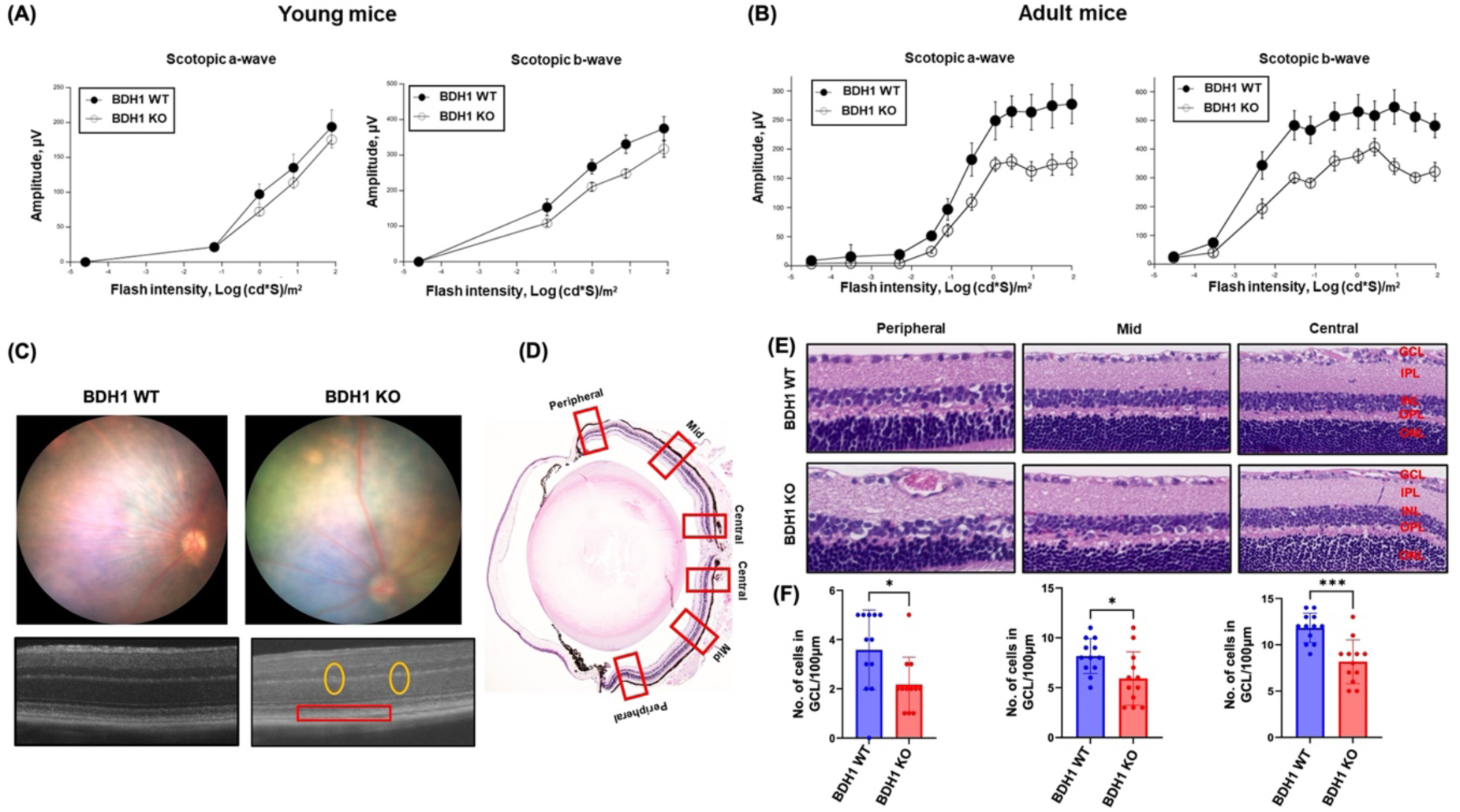
Global BDH1 deficiency impairs retinal structure and visual function. (A-B) Scotopic ERG recordings reveal reduced b-wave amplitudes in young BDH1 KO mice, with progressive declines in both a- and b-wave responses in older 6-8months old adult mice. (C) Fundus and OCT imaging demonstrate RPE irregularities (red box) and increased hyper-reflective foci (yellow circles) in BDH1 KO mice. (D) Representative H&E-stained retinal section of BDH1 KO mice showing regions (red blocks) used for cell counting. (E-F) BDH1 KO mice exhibit significantly reduced retinal ganglion cell (RGC) counts across the peripheral, mid, and central regions compared with WT. Data are mean ± SEM (n = 5–8 mice per group); *P≤0.05, **P≤0.001, ***P≤ 0.0001; ns, not significant.

Structural analyses by fundus photography and OCT imaging demonstrated RPE irregularities and increased hyper-reflective foci in BDH1 KO retinas (Figure 3C). The hyper-reflective foci observed in the INL of BDH1 KO mice likely indicate Müller glial activation or microglial infiltration, consistent with early degenerative remodeling (45, 46). Although overall retinal thickness remained unchanged (Suppl. Figure 1D), BDH1 KO mice displayed significant reductions in retinal ganglion cell (RGC) density (Figure. 3D-F), indicating neuronal vulnerability in BDH1 KO mice.

These results suggest that BHB is required for maintaining photoreceptor and inner retinal function. Given BHB’s role in mitochondrial redox balance and metabolism, BDH1 deficiency likely impairs critical cellular signaling and enhances oxidative stress in retinal neurons, leading to progressive functional decline. The observed RPE abnormalities and reduced ERG amplitudes in BDH1 KO mice are reminiscent of early pathological changes in models of oxidative or metabolic retinal degeneration (47–50), underscoring the essential role of BDH1 in retinal homeostasis and visual function.

To assess whether RPE-derived BHB is essential for retinal function, we generated mice with RPE-specific deletion of BDH1. RPE-specific BDH1 KO mice displayed a marked reduction of BDH1 expression in RPE lysates, while retinal lysates retained comparable expression levels to those of control mice, confirming RPE-specific BDH1 deletion. (Suppl. Figure 2A-B). However, RPE-specific BDH1 KO mice exhibited no alterations in ERG responses or retinal morphology as determined by fundus and OCT imaging compared to the control mice (Suppl. Figure 2C-E). These findings indicate that RPE-derived BHB is dispensable for retinal function when systemic, hepatocyte-derived BHB remains available. Future studies using hepatocyte-specific BDH1 KO mice will determine whether RPE-produced BHB becomes critical for retinal function under conditions of systemic BHB deficiency.

### Transcriptomic profiling reveals Müller cell dysfunction and retinal ganglion cell degeneration in BDH1-deficient retinas

Bulk RNA sequencing of BDH1 WT and KO ocular tissues revealed 1090 upregulated and 743 downregulated genes (raw p < 0.05; |log_2_FC| ≥ 0.25; Figure. 4A, volcano plot). Cell-type deconvolution analyses using an established retinal cell atlas suggested alterations in the composition of retinal cell types in BDH1-KO mice, with an apparent reduction in photoreceptors and RGCs, and an increase in microglial signatures (Figures 4B-C, Suppl. Figure 3A). Gene ontology and pathway enrichment analyses indicated activation of cytokine-mediated and tumor necrosis factor (TNF) signaling pathways, as well as microglial phagocytosis pathways. Conversely, downregulated pathways included genes involved in gliogenesis, inwardly rectifying potassium channels, voltage-gated calcium channels, glutamate metabolism, glutamatergic synaptic transmission, and long-term potentiation (Figure. 4D). We compared the expression of different retinal cell markers between BDH1-WT and KO ocular tissues (Suppl. figure 3B) as previously described in the literature (32–34). Microglial markers *Cd68, Ctss, Itgax, Itgb5*, and complement components *C1qa, C1qb, C1qc* were upregulated, consistent with a switch from a homeostatic surveillance state to an activated, complement-engaged phagocytic phenotype associated with synaptic pruning and neuroinflammation, a signature feature of retinal injury, glaucoma, and degenerative models (51–54). Conversely, Müller glial cell markers *Kcnj10, Sox2, Aqp4, Glul,* and *Slc1a3* were downregulated, suggesting impaired ionic buffering and glutamate clearance (55–57). RGC markers, Tubb3, fbxo44, Nell1, Slc17a6, Pou4f1, Rbpms2 were also reduced, consistent with compromised neuronal integrity (58). qRT-PCR validation confirmed downregulation of Müller glial cell genes *Scl1a3, Glul, Aqa4, and Kcnj10* in BDH1-KO retinas (Figure 4E-H), supporting potential defects in glial mediated extracellular glutamate and potassium buffering that can lead to impaired glutamate metabolism and recycling in the retina. Likewise, RGC markers, *Rbpms2, Nell1, Scl17a6, and Pouf4* were significantly decreased, implicating progressive neuronal degeneration (Figure 4I-L). Rod-specific markers, *GNAT1*, and *rhodopsin* (Suppl. Figure 1E-F) and cone-specific makers, *GNAT2, SWL-opsin, and MWL-opsin* (Suppl. Figure 1G-I) were also reduced in BDH1 KO retinas compared to WT retinas. Together, these data suggest that BDH1 deficiency disrupts Müller glial-neuronal coupling, impairs glutamate metabolism, promotes microglial activation, and is associated with progressive neurodegeneration.

**Figure 4:**
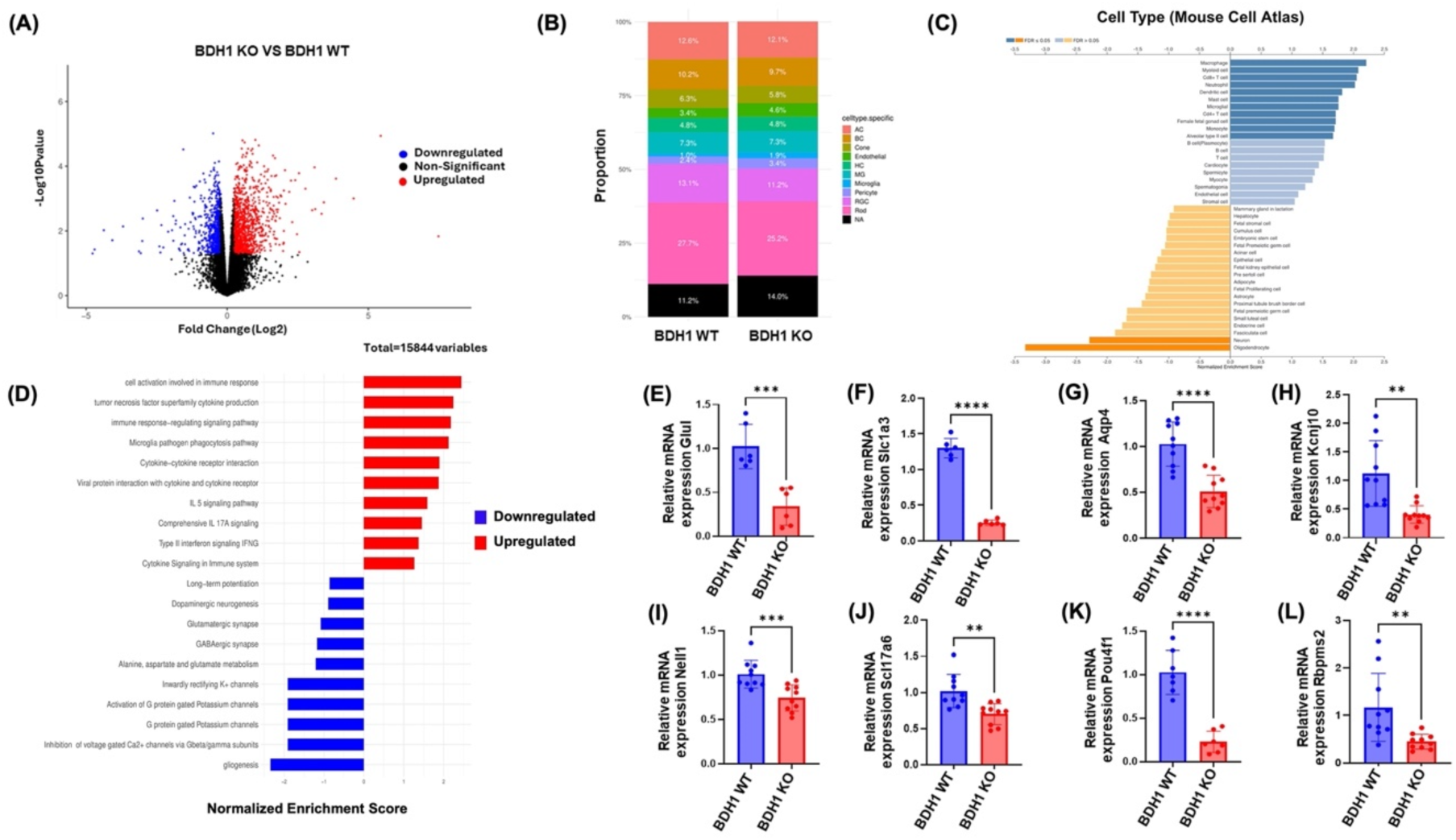
Transcriptomic profiling reveals cell-type and pathway alterations in BDH1-deficient ocular tissues. (A) Volcano plot showing differentially expressed genes in BDH1 KO versus WT retinas; blue and red indicate downregulated and upregulated genes in the BDH1 KO mice with a log2fold-change>0.25 and raw p value < 0.05. (B-C) Cell-type deconvolution analysis indicates reduced RGC and increased microglial cell numbers in BDH1 KO retinas. (D) Gene ontology (GO) enrichment highlights altered signaling cascades and biological processes upregulated or downregulated in BDH1 KO tissue. Quantitative RT-PCR validation shows decreased expression of (E-H) Müller cell-specific genes and (I-L) RGC-specific genes in BDH1 KO mice. Data represent mean ± SEM (n = 5-6 mice per group); *P≤0.05, **P≤0.001, ***P≤ 0.000; ns, not significant.

### BDH1 deficiency disrupts glutamate metabolism and induces oxidative stress

The downregulation of Müller cell-specific and RGC-specific genes in BDH1-KO retinas suggests a disruption of glial-neuronal metabolic coupling that is essential for retinal homeostasis (59). We therefore evaluated the expression of excitatory amino acid transporter (EAAT1), the primary glutamate transporter in Müller cells. EAAT1 expression was markedly reduced in BDH1 KO retinas, as demonstrated by immunoblotting (Figure 5A). The concurrent reduction in EAAT1 together with decreased glutamine synthetase (Glul) (Figure 4E), a key enzyme responsible for glutamate clearance and conversion to glutamine, suggests impaired retinal glutamate homeostasis. Such dysregulation may predispose the retina to excitotoxic stress and contribute to RGC vulnerability.

**Figure 5:**
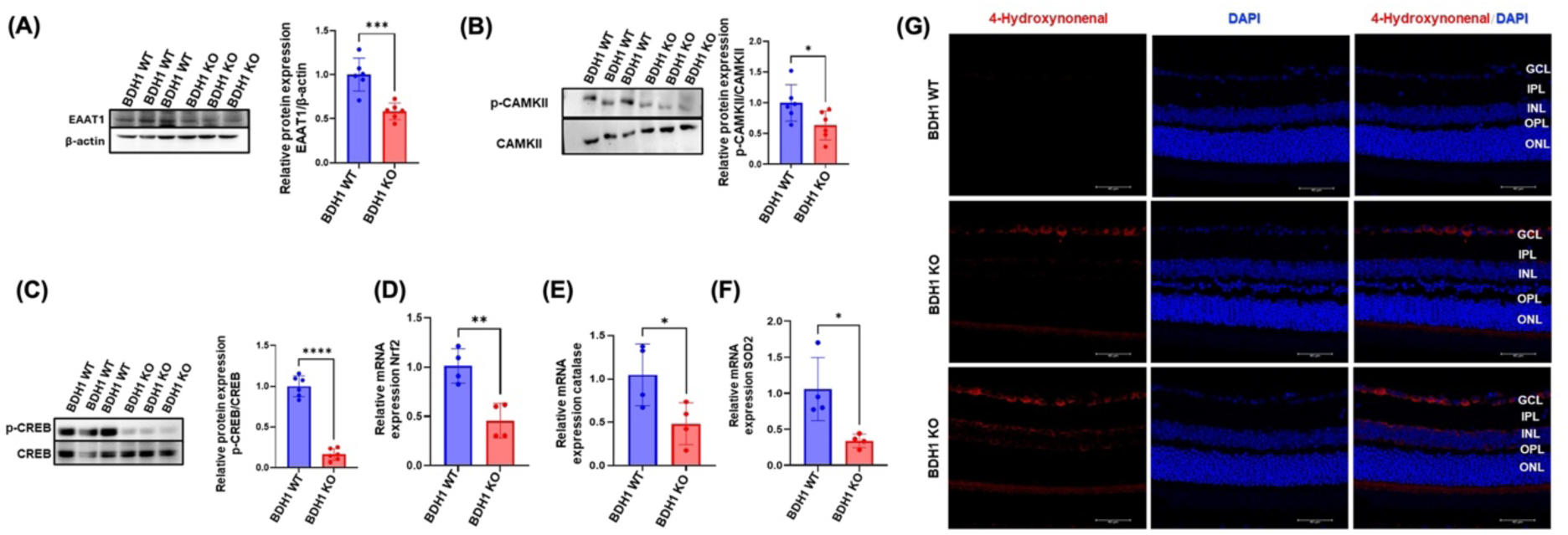
BDH1 deficiency impairs glutamate metabolism and induces oxidative stress. (A) Immunoblot analyses show decreased EAAT1 expression in BDH1 KO retinas. (B) p-CAMKII and (C) p-CREB levels are significantly decreased in BDH1 KO retinas compared with WT by immunoblotting analyses. (D-F) Quantitative RT-PCR demonstrates downregulation of *Nrf2* and its downstream antioxidant enzymes *SOD2* and *CAT* in BDH1 KO retinas. (G) BDH1 KO retinas show increased staining for oxidative stress and lipid peroxidation marker 4-hydroxynonenal compared with WT retinas. Data represent mean ± SEM (n = 5–6 mice per group); *P≤0.05, **P≤0.001, ***P≤ 0.0001; ns, not significant.

To investigate the molecular mechanisms underlying the reduced EAAT1 expression in BDH1 KO mice, we examined the expression of key signaling regulators. Both phosphorylated CREB (p-CREB) and phosphorylated CAMKII (p-CAMKII) were significantly decreased in BDH1 KO retinas compared to WT controls (Figure 5B-C). Since CREB activation drives *Slc1a3* transcription through calcium/CAMKII-dependent pathways (60, 61), reduced p-CREB and p-CAMKII levels indicate a suppression of the CAMKII-CREB transcriptional axis, potentially contributing to the downregulation of EAAT1 in BDH1 KO ocular tissues.

A marked decrease in *Nrf2* and its downstream antioxidant enzymes, superoxide dismutase 2 *(SOD2),* and catalase *(CAT)* (Figures 5D-F) was observed in BDH1 KO retinas, reflecting a weakened antioxidant defense in the retinal microenvironment. Additionally, retinal cross sections of BDH1 KO mice showed a significant increase in oxidative stress marker 4-hydroxynonenal (HNE), a lipid peroxidation product, in GCL, IPL, INL and to a lesser extent in photoreceptor inner segments (Figure 5G), indicating enhanced oxidative stress in the BDH1 KO retinas. Collectively, these findings suggest that BDH1 deficiency disrupts retinal metabolic and redox balance, characterized by impaired glutamate metabolism and reduced antioxidant capacity, thereby rendering the retina more vulnerable to stress-induced damage.

### BHB treatment restores CAMKII-CREB-EAAT1 signaling in BDH1 KO Müller cells

Primary Müller cells isolated from BDH1 WT and KO mice were first characterized for cellular purity using immunostaining for vimentin, a well-established Müller glial marker (Figure 6A). Further, quantitative real-time PCR confirmed the lack of BDH1 expression in the isolated primary Müller cells from BDH1 KO mice (Figure 6B).

**Figure 6:**
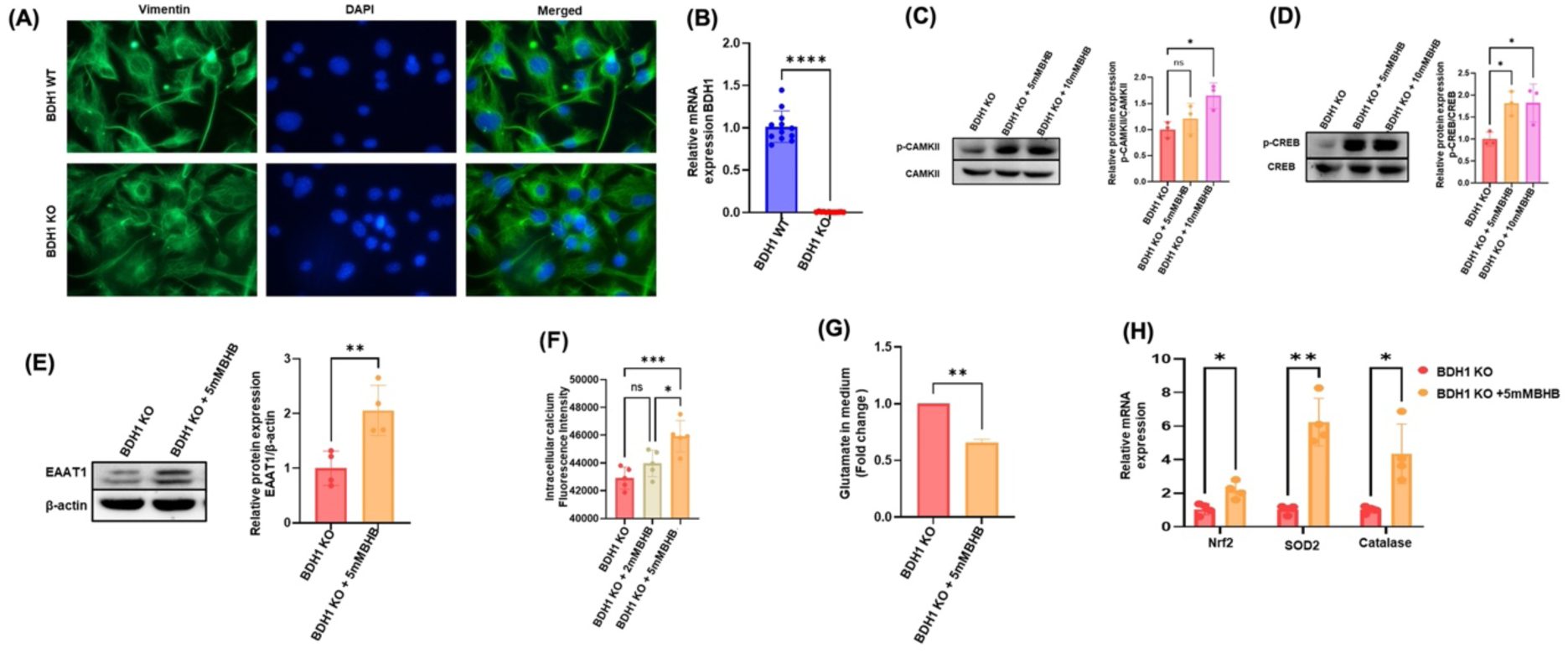
BHB treatment restores CAMKII-CREB-EAAT1 signaling in primary Müller cells. (A) Immunofluorescence staining for Vimentin confirms purity of isolated primary Müller cell cultures. (B) Real-time PCR confirms lack of BDH1 expression in cultured KO primary Müller cells. Immunoblot analyses show a significant increase in (C) p-CAMKII, (D) p-CREB, and (E) EAAT1 levels following BHB treatment compared to untreated control KO primary Müller cells. (F) BHB treatment increases intracellular calcium levels in BDH1 KO primary Müller cells. (G) Glutamate uptake assays demonstrate significantly improved glutamate clearance after BHB treatment in KO primary Müller cells. (H) Quantitative RT-PCR shows upregulation of *Nrf2* and its downstream antioxidant enzymes *SOD2* and *CAT* with BHB-treatment compared to untreated BDH1 KO Müller cells. Data represent mean ± SEM (n = 3); *P≤0.05, **P≤0.001, ***P≤ 0.0001; ns, not significant.

BDH1 KO primary Müller cells treated with BHB for 24 hours markedly restored p-CAMKII and p-CREB levels and upregulated EAAT1 expression compared to untreated BDH1 KO control cells (Figures 6C-E), indicating reactivation of the CAMKII-CREB signaling cascade known to drive Slc1a3 or EAAT1 transcription (60, 61). Functionally, BHB treatment significantly elevated intracellular calcium levels in Müller cells, consistent with enhanced calcium signaling required for CREB activation (Figure 6F). This was corroborated by glutamate uptake assays, which showed substantial improvement in glutamate clearance following BHB treatment compared with untreated KO controls (Figure 6G).

Furthermore, BHB treatment restored the expression of *Nrf2, SOD2, and CAT,* reflecting rejuvenation of the antioxidant defense system and reduced oxidative susceptibility (Figure 6H). Together, these findings demonstrate that BHB treatment effectively restores Müller cell functional integrity by reactivating the CAMKII-CREB-EAAT1 pathway and enhancing both antioxidant capacity and glutamate homeostasis.

### Exogenous BHB treatment restores retinal structure and CREB-EAAT1 signaling in BDH1-deficient mice

To determine whether BHB supplementation could rescue retinal defects *in vivo*, BDH1 KO mice were administered exogenous BHB (100mg/kg bodyweight) for 15 days. As expected, there was a significant increase in the serum BHB levels in the BDH1 KO-BHB treated group compared to the untreated BDH1 KO group (Figure 7A).

**Figure 7:**
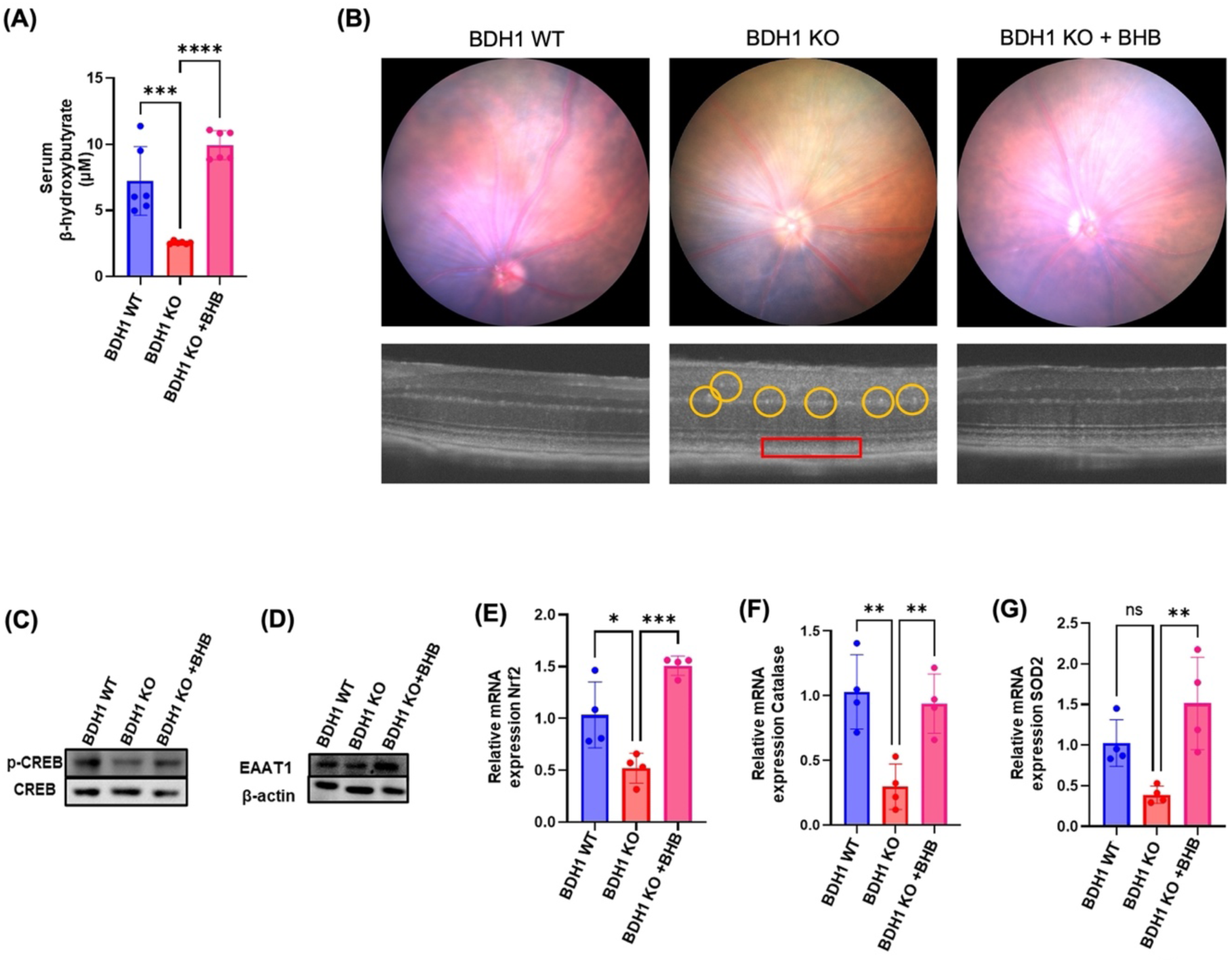
Exogenous treatment of BHB restores retinal structure and reactivates CREB-EAAT1 signaling in BDH1 KO mice. (A) Serum BHB levels are increased in BHB-treated BDH1 KO mice compared to the untreated group (B) Fundus and OCT imaging show reduced RPE irregularities (red box) and hyper-reflective foci (yellow circles) in BHB-treated BDH1 KO compared to untreated BDH1 KO retinas. (C-D) Immunoblot analyses reveal a significant increase in p-CREB and EAAT1 expression in BHB-treated BDH1 KO retinas compared to untreated BDH1 KO retinas. (E-G) Quantitative RT-PCR confirms upregulation of *Nrf2* and its downstream antioxidant enzymes *SOD2* and *CAT* in BHB-treated BDH1 KO retinas compared to untreated BDH1 KO retinas. Data represent mean ± SEM (n = 5-6 mice per group); *P≤0.05, **P≤0.001, ***P≤ 0.0001; ns, not significant.

Fundus and OCT imaging revealed reduced RPE irregularities and hyper-reflective foci (Figure 7B), while retinal lysates exhibited a significant increase in p-CREB and EAAT1 expression (Figures 7C-D) in BHB-treated BDH1 KO mice compared to untreated BDH1 KO mice. Quantitative RT-PCR demonstrated a robust upregulation of *Nrf2* and its key downstream antioxidant genes *SOD2* and *CAT* in the BHB-treated KO group (Figure 7E-G), suggesting restored redox homeostasis.

Together, these findings indicate that exogenous BHB supplementation reinstates the CREB-EAAT1 signaling axis, augments antioxidant capacity, and improves retinal structural integrity, effectively reversing metabolic and oxidative dysregulation associated with BDH1 deficiency.

## Discussion

The neurosensory retina exhibits exceptionally high metabolic activity and depends on tightly coordinated metabolic exchange among the RPE, photoreceptors, Müller glia, and RGCs. This reliance renders the retina particularly vulnerable to mitochondrial dysfunction, redox imbalance, and impaired metabolic coupling, which are critical processes implicated in AMD and other degenerative retinopathies (62). Retinal aging and associated disorders such as AMD, DR, and glaucoma arise, in part, from cumulative oxidative stress, chronic inflammation and progressive bioenergetic decline that converge on neuronal degeneration (63).

Although KBs are typically viewed as alternative energy substrates during fasting, growing evidence indicates that they also serve as constitutive regulators of redox balance, transcription pathways, and intracellular signaling (1). Given the retina’s high metabolic demand and the intrinsic ketogenic capacity of the RPE (7, 8), KBs may contribute to retinal homeostasis beyond intermittent energy supplementation. However, the role of basal ketone metabolism in retinal physiology has remained largely unexplored.

In human donor eyes with AMD, we observed a marked reduction in BDH1 expression across multiple retinal layers. This observation is notable because hepatic BDH1 expression and circulating ketone levels decline with age in mice (40), suggesting that impaired ketone metabolism may constitute an intrinsic component of AMD pathophysiology. Also, these findings align with extensive evidence linking mitochondrial dysfunction and age-related metabolic decline to retinal degeneration (63). The antioxidant, anti-inflammatory and epigenetic functions of BHB (10–12) suggest that ketone-dependent pathways may mitigate key pathological features of retinal degeneration including oxidative stress, inflammasome activation, microglial reactivity and synaptic vulnerability. Consistent with this concept, feeding ketogenic diet in glaucoma models enhances retinal oxidative metabolism, increases MCT expression, improves NAD+/NADH ratios, and preserves RGC soma and axons (64). Mechanistically, BHB confers neuroprotection in part through activation of Nrf2 dependent antioxidant pathways (17, 65, 66), and inhibition of the NLRP3 inflammasome signaling (67, 68).

Global BDH1 knockout mice recapitulated many features of retinal dysfunction. Young BDH1-deficient mice exhibited selective reductions in scotopic and photopic b-wave amplitudes, indicating inner retinal impairment involving ON-bipolar cells, Müller glia or synaptic transmission. With aging, these abnormalities progressed to reductions in both a- and b-waves, suggesting combined photoreceptor and inner retinal degeneration. Systemic metabolic parameters, including glucose tolerance, serum triglycerides, and body weight remained unaltered, indicating that retinal dysfunction resulted from intrinsic defects in ketone metabolism rather than systemic metabolic derangements. These findings are in concordance with studies on mitochondrial pyruvate carrier-1 (MPC1)-deficient mice, where metabolic impairment disrupts retinal signaling before structural degeneration becomes apparent (69).

Transcriptomic profiling revealed pronounced Müller glial dysfunction and robust activation of innate immune and microglial pathways in BDH1-deficient retinas. Müller cells regulate synaptic transmission, potassium buffering, neurotransmitter recycling and antioxidant defense (70). Consistent with these functions, BDH1-deficient retinas exhibited marked reductions in the expression of *Kcnj10, Aqp4, Glul, and Slc1a3* genes essential for glutamate clearance and synaptic homeostasis (71, 72). Loss of *Slc1a3/*EAAT1, as demonstrated in EAAT1-null mice, elevates extracellular glutamate, diminishes b-wave amplitudes and promotes RGC degeneration (71). Thus, early b-wave reductions observed in BDH1-deficient mice likely reflect impaired Müller cell-mediated support. Concomitantly, BDH1 loss induced microglial activation and complement cascade upregulation, including increased expression of *Ctss, Itgax and C1q* expression. Complement-dependent synaptic opsonization and microglial engulfment are central mechanisms of RGC synaptic loss in multiple retinopathies (73). Together, impaired Müller cell support and microglial activation create a feed forward neuroinflammatory milieu that exacerbates synaptic injury and RGC vulnerability, consistent with the progressive degeneration observed histologically and functionally in BDH1-deficient mice.

Mechanistically, BDH1 deficiency disrupted glutamate homeostasis through impaired EAAT1 expression in Müller cells. The glutamate transporter EAAT1, predominantly expressed in Müller glia, is the primary transporter responsible for reuptake of excess glutamate from the synaptic cleft, thereby maintaining normal neurotransmission and preventing excitotoxicity (60). Reduced EAAT1 increases extracellular glutamate in the synapses, driving calcium-dependent excitotoxic injury and neuronal loss (74, 75). We found that BDH1-deficient retinas exhibited reduced phosphorylation of CAMKII and CREB, transcriptional regulators of EAAT1. Prior work in brain astrocytes demonstrates that exogenous BHB treatment increases intracellular Ca^2+^, activates CAMKII, enhances CREB phosphorylation, and induces EAAT1 transcription (63). Consistent with this pathway, exogenous BHB supplementation restored CAMKII-CREB signaling in primary Müller cells and *in vivo* in BDH1 KO retinas, accompanied by increased EAAT1 expression and enhanced glutamate uptake. These results define a BDH1-dependent CAMKII-CREB-EAAT1 signaling axis that maintains constitutive glutamate homeostasis in retina.

Beyond neurotransmission, Müller cells support retinal antioxidant defenses through *Nrf2*-dependent detoxification pathways and glutathione synthesis (76, 77). BDH1 deficiency reduced retinal *Nrf2, SOD2 and CAT* expression and increased the oxidative stress and lipid peroxidation marker 4-hydroxynonenal, indicating diminished antioxidant capacity and exacerbated oxidative stress in BDH1 KO retinas. BHB supplementation restored these antioxidant pathways in BDH1-deficient Müller cells and BDH1 KO retinas. Since Müller cell dysfunction in neurotransmission and redox balance is strongly implicated in AMD and related retinopathies (78), BDH1 loss likely contributes to neuronal injury through combined synaptic dysfunction and redox imbalances.

With aging, BDH1-deficient mice developed ERG a-wave deficits, photoreceptor dysfunction, cone degeneration and RGC loss, indicating progression to pan-retinal pathology. Photoreceptors rely on systemic- and RPE-derived metabolites for outer segment renewal and redox homeostasis. Impaired ketone oxidation therefore compromises energy balance, increases ROS, and destabilizes RPE-photoreceptor interactions (7). These alterations resemble metabolic features of early AMD, in which mitochondrial stress and impaired metabolite exchange precede photoreceptor degeneration (63). Importantly, exogenous BHB supplementation restored Müller cell homeostasis, attenuated inflammatory and oxidative stress signatures, improved glutamate clearance, and ameliorated structural deficits in BDH1-deficient mice. These findings suggest that therapeutic enhancement of ketone signaling, through BHB administration, ketogenic diet, or pharmacologic modulation, may support retinal metabolism and slow neurodegeneration.

Despite establishing BDH1 as an essential metabolic regulator in the retina, several questions remain. The upstream mechanisms driving BDH1 downregulation in AMD require further investigation. Moreover, RPE-specific BDH1 deletion did not replicate the retinal pathology observed in global BDH1 KO mice, suggesting that systemic ketone availability contributes to retinal integrity. Future studies employing hepatocyte-specific, Müller cell–specific, and photoreceptor-specific BDH1 deletion models will be critical for delineating the relative contributions of systemic versus local ketone metabolism in maintaining retinal health. Additionally, although BHB treatment restored key protective pathways, therapeutic dosing, delivery strategies, and long-term safety require further evaluation.

In summary, our findings identify BDH1-dependent ketone metabolism as a constitutive requirement for retinal homeostasis. Loss of BDH1 disrupts Müller cell function, perturbs glutamate and redox balance, activates microglia, and drives progressive neuronal degeneration, establishing the BDH1-BHB axis as a central determinant of retinal resilience (Figure 8). Restoration of molecular and structural integrity by exogenous BHB in BDH1-deficient retinas highlights the therapeutic potential of targeting ketone metabolism to attenuate retinal degeneration, including AMD, where BDH1 is intrinsically reduced.

**Figure 8:**
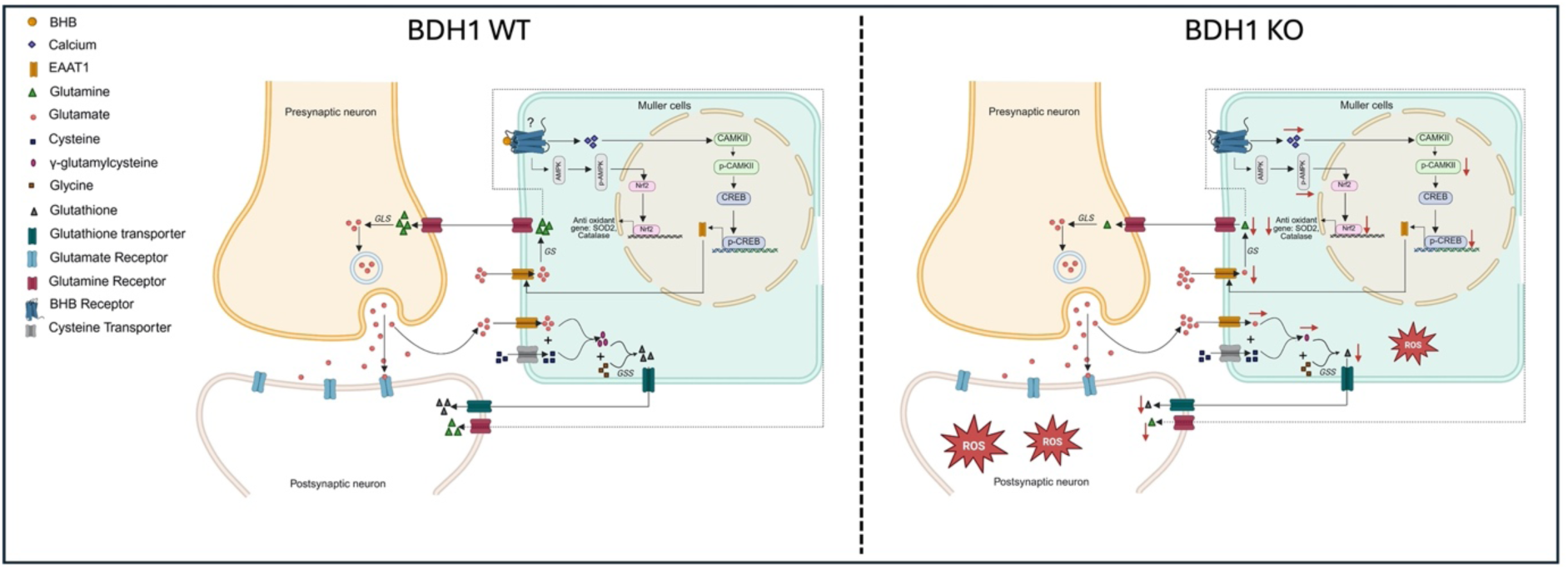
Schematic of BHB-mediated interactions between Müller glial cells and pre-and post-synaptic neurons. The left panel illustrates signaling in BDH1 WT retinas, where BHB modulates intracellular signaling pathways, activating the CAMKII-CREB axis and leading to increased *Slc1a3* expression. Elevated *Slc1a3* or EAAT1 expression enhances glutamate uptake from the synaptic cleft, thereby limiting glutamate-induced excitotoxicity. Once transported into the Müller glial cells, glutamate is converted to glutamine by glutamine synthetase (GS), and is returned primarily to presynaptic neurons for resynthesis of glutamate via glutaminase (GLS). Additionally, glutamate also contributes to glutathione synthesis through sequential reactions catalyzed by glutamate-cysteine ligase (GCL) and glutathione synthetase (GSS). Müller cells supply glutathione and its precursors, supporting neuronal antioxidant defenses. BHB can activate signaling pathways like AMPK that enhance Nrf2-dependent antioxidant responses, and increase enzymes such as SOD2 and CAT, thereby mitigating oxidative stress. The right panel depicts how impaired BHB metabolism in BDH1 KO retinas may contribute to inadequate glutamate clearance and increased oxidative stress in Müller glia and neurons.

## Author Contributions

R.G. and J.P.G-P. designed the experiments. R.G., E.K., D.D., S.P., E.N., M.B., N.P., and O.K. performed the experiments and data analysis. R.G, D.D., E.N., M.B., N.P., O.K., J.P.G-P. wrote and edited the manuscript. Funding and supervision by J.P.G-P. All authors have read and agreed to the final version of the manuscript.

## Acknowledgement

We thank Dr. Henry J. Kaplan at Saint Louis University for his assistance with the procurement of human AMD samples and for valuable input on fundus and OCT imaging analyses.

