## Supplementary Figures for "BDH1-Dependent Ketone Body Metabolism Maintains Müller Cell Homeostasis and Retinal Function"

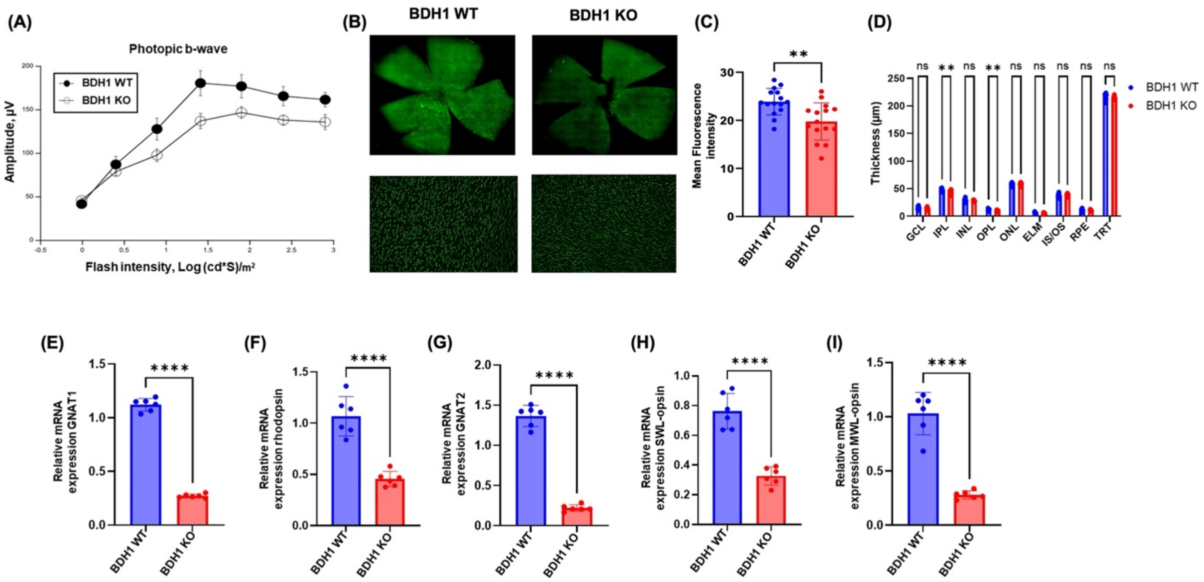


Supplementary Figure 1

**Supplementary Figure 1: Global BDH1 deficiency reduces cone function and density without altering retinal thickness.** (A) Photopic ERG shows reduced b-wave amplitudes in young BDH1 KO mice. (B) Representative image of cones labeled with FITC-PNA in flat mount retinas and (C) quantification of the mean fluorescence intensity reveals decreased cone density in BDH1 KO mice. (D) Quantification of total retinal thickness (TRT) shows no genotype-dependent differences. (E-F) Real-time PCR shows reduced rod-specific gene expression in BDH1 KO retinas. (G-I) Real-time PCR shows reduced cone-specific gene expression in BDH1 KO compared to BDH1 WT retinas. Data represent mean ± SEM (n = 5-6 mice per group); *P≤0.05, **P≤0.001, ***P≤ 0.0001; ns-not significant. Abbreviations: Ganglion cell layer (GCL), inner plexiform layer (IPL), inner nuclear layer (INL), outer plexiform layer (OPL), outer nuclear layer (ONL), external limiting membrane (ELM), inner segment/outer segment (IS/OS), retinal pigment epithelium (RPE) and total retinal thickness (TRT).

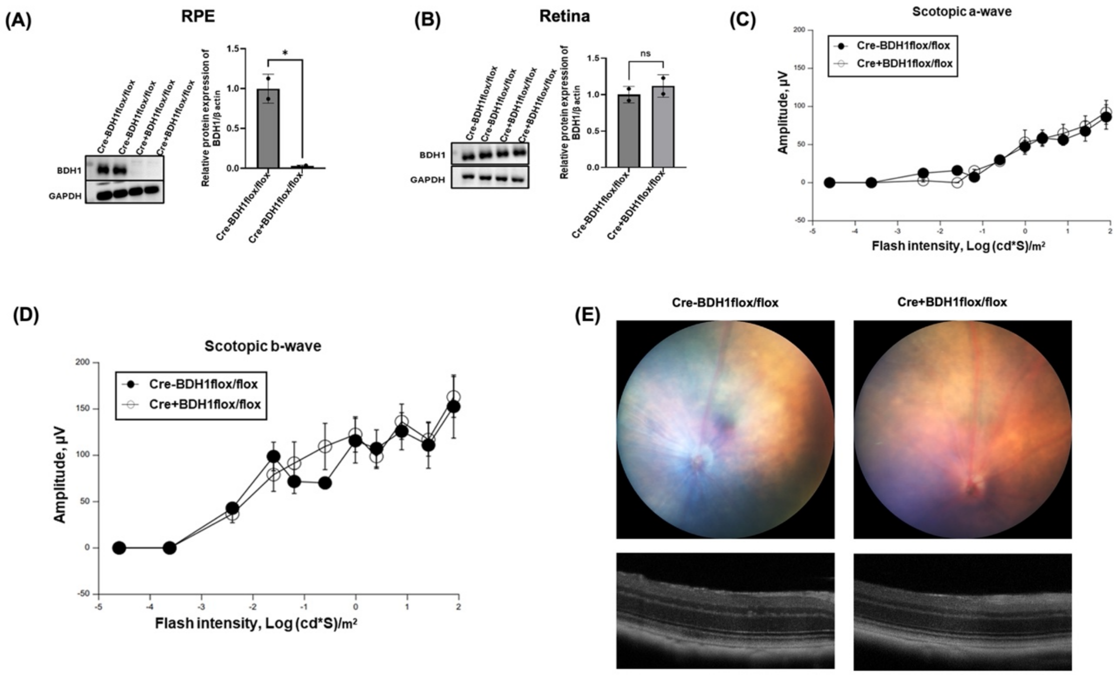


Supplementary Figure 2

**Supplementary Figure 2: RPE-specific conditional BDH1 deletion preserves retinal structure and function.** Immunoblot analyses confirms (A) complete lack of BDH1 protein in RPE lysates but (B) no significant change in retinal lysates from RPE-specific BDH1 KO mice ($Cre^{+}Bdh1^{fl/fl}$) compared to Cre- control mice. (C-D) Scotopic ERG responses show no significant differences between genotypes. (E) Fundus and OCT imaging reveal no structural differences between Cre- and Cre+ mice. Data represent mean ± SEM (n = 5-6 mice per group); *P≤0.05, **P≤0.001, ***P≤ 0.0001; ns- not significant.

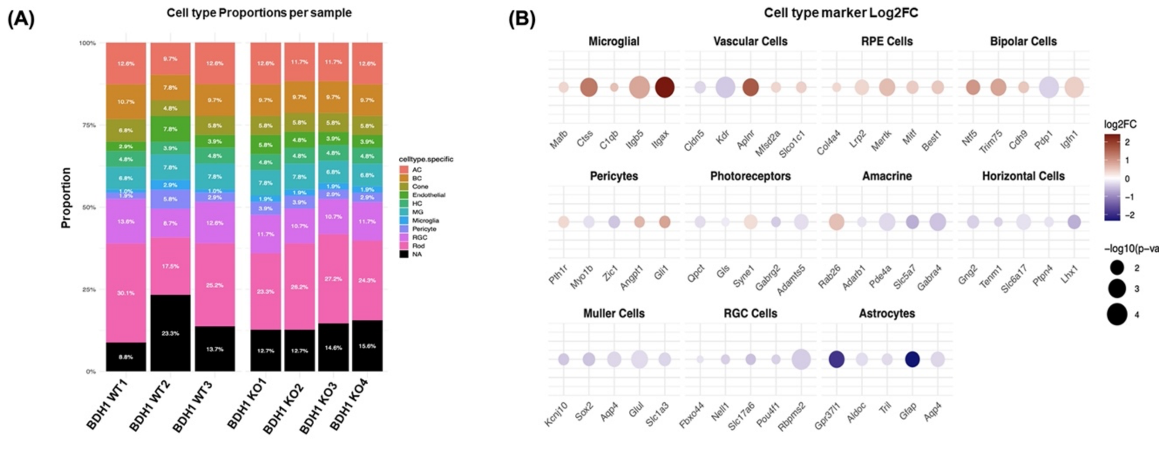


Supplementary Figure 3

**Supplementary Figure 3: Transcriptomic analysis of retinal cell type-specific markers** (A) Cell-type deconvolution analysis showing alterations in the composition of retinal cell types in BDH1-KO mice. (B) Differential expression analysis of retinal cell-type markers in BDH1 WT and KO mice.
